# Regulation of prostate androgens by megalin and 25-hydroxyvitamin D status: Mechanism for high prostate androgen in African American men

**DOI:** 10.1101/2021.11.09.467567

**Authors:** Jason Garcia, Kirsten D. Krieger, Candice Loitz, Lillian Perez, Zachary A. Richards, Yves Helou, Steve Kregel, Sasha Celada, Clementina A. Mesaros, Peter H. Gann, Thomas E. Willnow, Donald Vander Griend, Rick Kittles, Gail S. Prins, Trevor Penning, Larisa Nonn

## Abstract

Vitamin D deficiency is associated with an increased risk of prostate cancer (PCa) mortality and is hypothesized to contribute to PCa aggressiveness and disparities in African American populations. The prostate epithelium was recently shown to express megalin, an endocytic receptor that internalizes globulin-bound hormones, which suggests regulation of prostate hormone levels, in contrast to the free hormone hypothesis. Here, we demonstrated that megalin imports testosterone bound to sex hormone-binding globulin into prostate cells. Prostatic loss of *Lrp2* (megalin) in a mouse model resulted in reduced prostate testosterone and dihydrotestosterone (DHT) levels. Megalin expression was regulated and suppressed by 25-hydroxyvitamin D (25D) in cell lines, patient-derived prostate epithelial cells, and prostate tissue explants, indicating a negative feedback loop. In patent samples, the relationships between hormones support this feedback mechanism, as prostatic DHT levels are higher in African American men and are inversely correlated with serum 25D status. Megalin levels are reduced in localized PCa by the Gleason grade and in patients with future disease recurrence. Our findings suggest that the free hormone hypothesis should be revisited for testosterone and highlight the impact of vitamin D deficiency on prostate androgen levels, which are known drivers of PCa. Thus, we revealed a mechanistic link between vitamin D and PCa disparities observed in African Americans.

## INTRODUCTION

Prostate cancer (PCa) is the second most frequently diagnosed cancer in men in the United States, and African American (AA) men have 60% higher incidence and 200% higher mortality rates than European American (EA) men ^1^. The reason for this disparity is multifactorial and involves both biological and socioeconomic factors; however, when controlled for, AA men still present with more aggressive disease at a younger age^2^. AA men are also at increased risk of vitamin D deficiency^3^ due to melanin, which decreases cutaneous synthesis of cholecalciferol (pre-vitamin D) from 7-dehydrocholesterol. Cholecalciferol is converted by the liver to the main circulating form of vitamin D, 25-hydroxyvitamin D (25D), which is the prohormone for 1,25-dihydroxyvitamin D (1,25D), a steroid hormone essential for normal human physiology and calcium homeostasis ^4^. Thus, vitamin D status is dependent on vitamin D supplementation, sun exposure, and skin pigmentation^3^.

Circulating levels of 25D are inversely correlated with aggressive PCa ^5–10^ and numerous anti-cancer activities have been demonstrated for vitamin D metabolites in vitro and in vivo^9,11,12^. Thus, differences in vitamin D status are hypothesized to contribute to the disparity in PCa incidence in AA men.

25D and other hormones are thought to follow the free hormone hypothesis which posits that that intracellular concentrations of hormones are dependent on passive diffusion of “free” hormones not bound by serum globulins ^13^. However, we demonstrated that circulating and prostate tissue levels of vitamin D metabolites did not correlate, indicating that passive diffusion of unbound hormones does not driving prostate concentrations ^14^. Consistent with a hormone import mechanism, we further demonstrated that prostate epithelial cells express megalin, an endocytic receptor encoded by *LRP2*, with a well-characterized function of binding and internalizing globulin-bound 25D from the glomerular filtrate ^15,16^. We further showed that prostatic expression of the megalin gene (*LRP2*) was negatively correlated with 25D levels only in AA men^14^. *LRP2* was also positively correlated with the percentage of West African ancestry in the cohort^14^. These findings suggest that the free hormone hypothesis may not apply to the prostate and that a compensatory mechanism may increase prostate megalin levels when systemic levels of 25D are deficient.

Megalin binds to and internalizes sex hormone-binding globulin (SHBG), a serum transporter of testosterone (T). Megalin import of SHBG-bound T occurs most notabley in kidney cells, but import of SHBG has also been shown in LNCaP PCa cells ^17^. Circulating T concentrations were not correlated with intraprostatic concentrations, further supporting an alternative to passive diffusion ^18^. Megalin-deficient mice exhibit defects in the maturation of their reproductive organs, suggesting dysregulation of sex hormones^19^. Polymorphisms in the megalin gene *LRP2* are associated with PCa recurrence, PCa-specific mortality, and the effectiveness of androgen deprivation therapy (ADT)^20^.

Here, we examined megalin as a mechanism to increase prostatic hormone import and the regulation of megalin by vitamin D metabolites. This mechanism is highly relevant to PCa disparities, given that androgens contribute to PCa pathogenesis and AA men are more likely to be 25D deficient. Here, we show the mechanistic examination of steroid hormone transport by megalin in prostate cells, a knockout mouse model, and patient prostate tissue explants. We further validated the findings in clinical prostate specimens.

## MATERIALS AND METHODS

### Patient sera and prostate tissue

Samples were de-identified and collected from the three cohorts. Cohort 1 included 60 patients:30 from the UIC Hospital (Chicago, Illinois, USA) and 30 from the Cooperative Human Tissue Network (CHTN) Western Division at Vanderbilt University (Nashville, Tennessee, USA). The CHTN samples were acquired under exemption (IRB 2018-0281). UIC fresh-frozen prostate samples were collected from radical prostatectomy patients under IRB 2004-0679 and 2015-1294. UIC serum was collected from patients under IRB 2017-0807. All the patients had localized PCa without prior chemotherapy or hormone therapy. Cohort 2 included only prostate tissue from UIC patients collected under IRB 2004-0679. Cohort 3 comprised serum only and was acquired the Prostate Cancer Biorepository Network (PCBN). Because the samples were de-identified, the research was determined to not fit the definition of human subjects research by the UIC IRB (2013-0341).

### Hormone measurement in patient samples

#### Calibration curve

Standard compounds testosterone (T) and dihydrotestosterone (DHT) and the internal standard (IS) d3T were purchased from Cerilliant (Round Rock, TX, USA). Nine calibrators (0.0625, 0.125, 0.25, 0.5, 1, 5, 10, 50, and 100 ng/mL in methanol) were used to establish calibration curves with the spiked-in IS. The curves were fitted by linear regression with a weighting factor of 1/*x*.

#### Sample preparation and extraction

Tissue samples and ISs were mixed and bead homogenized using a Mikro-Dismembrator II (Handelskontor Freitag, Germany) prior to extraction. Extraction was performed three times with hexane:ethyl acetate (60:40 [vol:vol]). The organic layer from each extraction was collected, combined, and dried under nitrogen. The residue was reconstituted in methanol:water (20:80 [vol:vol]) and subjected to solid-phase extraction using an ISOLUTE C18 SPE cartridge (100 mg, 1 mL), following the manufacturer’s protocol. The final eluate was dried before LC–MS analysis. Human serum samples were extracted using the same procedure.

#### LC-MS/MS analysis

Quantification of T and DHT was performed using an SCIEX Qtrap 6500 spectrometer coupled with an Agilent 1290 ultra-performance liquid chromatography (UPLC) system. The dried sample was reconstituted in methanol and resolved using a Waters ACQUITY UPLC BEH C18 column (1.7 μm, 2.1 × 100 mm) maintained at 45°C at a flow rate of 450 μL/min. The elution started with 60% mobile phase A (5% methanol in water, 0.1% formic acid), followed by a linear gradient increase in mobile phase B (acetonitrile with 0.1% formic acid) from 40 to 80%. MS data were acquired by multiple reaction monitoring (MRM) in positive mode with an electrospray ionization (ESI) source voltage of 5.0 kV and a temperature of 450°C. T, DHT, and D3T were detected by monitoring their transitions to the signature product ions 289>97 (T), 291>255 (DHT), and 292>97 (D3T), respectively. Data were analyzed using the Analyst software.

#### Measurement of Vitamin D Metabolites

The extraction and measurement of 25D was previously reported by our group and was done on 50 μL of serum at Heartland Assays (Ames, IA, USA) ^21^.

### Cell lines

HEK293, LNCaP, and 22Rv1 cells were purchased from ATCC (VA, USA), and 957E-hTERT cells were generously donated by John Isaacs and maintained in keratinocyte serum-free medium (KSFM) (ThermoFisher Scientific). Cell lines were authenticated by short-tandem-repeat analyses prior to use. PrE-AA1 and PrE-AA2 are primary patient-derived epithelial cells derived in our lab using previously reported methods ^12,22^. HEK293 cells were maintained in Dulbecco’s Modified Eagle’s medium (DMEM) (Gibco) supplemented with 10% (v/v) fetal bovine serum (FBS) (Gibco). LNCaP and 22Rv1 cells were maintained in phenol-free Roswell Park Memorial Institute (RPMI) medium (Gibco) supplemented with 10% (v/v) FBS. Cells were switched to 5% (v/v) charcoal-stripped FBS (Millipore-Sigma) overnight and serum-starved for 1 h prior to experimentation. Primary prostate cells established from fresh male radical prostatectomy tissues were isolated as previously described and cultured in prostate cell growth medium (Lonza). All cells were cultured at 37°C and 5% CO_2_. All the cells are described in **Table S1**.

### RNA isolation and reverse transcription–quantitative PCR (RT–qPCR)

RNA was isolated using TRIzol reagent (Thermo Fisher Scientific). RNA concentration and quality were determined by measuring the absorbance ratio at 260/280 nm using a NanoDrop One spectrophotometer (Thermo Fisher Scientific). Total RNA (500 ng) was reverse-transcribed using the High-Capacity cDNA Reverse Transcription Kit (Applied Biosystems). The resulting cDNA was used for quantitative PCR amplification on a QuantStudio6 machine (Thermo Fisher Scientific) using gene-specific primers (**Table S2**) and FastStart Universal SYBR Green master mix (Millipore-Sigma). Reactions were run in duplicate and relative *C*_t_ values were normalized and calculated independently using the –ΔΔ *C*_t_ method for the expression of the housekeeping genes *HPRT1* and *RPL13A* (all primers are listed in **Table S1**).

### Western blotting

Cells were grown to 80% confluence, and protein lysates were collected in cell lysis buffer (9803, Cell Signaling Technology). Protein (30 μg) was electrophoresed on a Bis-Tris protein gel (NuPAGE) and transferred to a PVDF membrane. Membranes were blocked for 1 h using Odyssey Blocking Buffer (LiCOR) and probed with anti-megalin rabbit monoclonal antibody (1:1,000; M02463, Boster Bio) and anti-actin (1:1,000; 4499S, Cell Signaling Technology), and with secondary antibodies against rabbit and mouse (926-68071, LiCOR). Blots were imaged using an Odyssey CLx imaging system (LiCOR).

### T and SHBG treatments

LNCAP and 22Rv1 cells at 80% confluency were incubated with 25 nM T ± 125 nM human SHBG (SHBG-8259H, Creative BioMart) and 1 μM receptor-associated protein (MEG-Inh) (BML-SE552-0100, Enzo Life Sciences) for 16 h. T and SHBG were preincubated for 30 min before addition to cells. Cells were pretreated with megalin inhibitor for 1 h before hormone addition.

### Luciferase reporter assays

Cells at 70% confluence were transfected with luciferase plasmids, and luciferase activity was measured after 48 h using the dual-luciferase reporter assay system and GloMax 20/20 (Promega). pRL-null *Renilla* plasmid was cotransfected at 0.4 pg/μL to control for transfection efficiency. For *LRP2* promoter, 0.2 ng/μL *LRP2* promoter-driven *Renilla* luciferase reporter (S712992, Switchgear Genomics) and 0.4 pg/μL PGL4.50 *Photinus pyralis* (E310, Promega) luciferase reporter were simultaneously treated with 10 nM 25D or 10 nM T. *Renilla* luciferase relative light units are defined as the ratio of *Renilla* to *Photinus* activity.

### DBP-488 and SHBG-555 internalization

Recombinant human SHBG (ProSpec, Israel) was directly labeled with Alexa Fluor-555 using a protein conjugation kit (Thermo Fisher Scientific) according to the manufacturer’s protocol. Aliquots of the globulin conjugate were stored at −20°C until use. Cells were grown to 70% confluence in eight-well chamber slides and incubated with SHBG-555 alone, SHBG-555 +T, or MEG-Inh + SHBG-555 + T, as described above. After 4 h, the cells were counterstained with Alexa Fluor 647 phalloidin (F-actin) and DAPI (Thermo Fisher Scientific), and visualized by confocal microscopy.

### *Lrp2*-flox/Pb-MerCreMer mice

All procedures involving animals in this study were approved by the University of Illinois at the Chicago Office of Animal Care and Institutional Biosafety (OACIB) approved all procedures involving animals in this study. Transgenic mice harboring the probasin promoter driving MerCreMer were acquired from Jackson Laboratory (ProbasinBAC-MerCreMer or C57BL/6-Tg(Pbsn-cre/Esr1*)14Abch/J, strain 020287). Generation of mice with *loxP* sites flanking *Lrp2* exons 71 through 75 (*Lrp2*^fl/fl^) has been previously described ^23^. *Lrp2*^fl/fl^ mice were crossbred with homozygous Pb-MerCreMer (*Cre* ^+/+^) mice. F1 cross progenies were mated to generate *Lrp2*^fl/fl^/*cre*^+/+^ mice. Mice were genotyped using Transnetyx and injected with tamoxifen (TAM)(50 mg/kg) at two stages of development (P10 or 5 weeks). Control TAM-injected mice were *Lrp2*^fl/fl^ or *Cre* ^+/+^. To confirm recombination, DNA was isolated from tail snips and prostate cell pellets using a DNeasy Blood & Tissue kit (Qiagen), followed by PCR with DreamTaq Green PCR master mix (Thermo Fisher Scientific) using primers spanning exons 71–75 and primers spanning exons 76–77 as a control. PCR products were imaged using agarose gels. The primers used are listed in **Table S2**.

### Mouse DHT and T quantitation

A protocol similar to that described by Higashi et al. was followed ^24^. Briefly, the internal standard mix (500 pg each of T-IS, DHT-IS, E1-IS, and E2-IS and 100 pg of 3α- and 3β-diol) were added to 0.6 mL of 0.1 mM PBS in a homogenization vial kept in ice. Frozen tissue (approximately 20 mg) was cut on a tile in dry ice with a blade kept in dry ice and added directly to the homogenization vial. The tissue was homogenized twice for 10 min on ice in a bullet blender. The homogenate was transferred to a borosilicate tube and ethyl ether (4 mL) was added and shaken for 30 min, followed by incubation for 2 h at 50°C with shaking at 4°C overnight. The sample was then centrifuged at 1,500*g* for 10 min. The upper organic phase was transferred to a new borosilicate tube using a glass pipette. The organic phase was then dried under nitrogen. Samples were stored at −20°C before derivatization and LC-MS analysis, as previously described.

### Ex vivo prostate tissue slice culture

Fresh prostate tissue was obtained from radical prostatectomy patients at UIC with informed consent (IRB 2007-0694). Tissue from a 5-mm punch was sliced into 300-μm sections using an Alabama Tissue Slicer (Alabama Research and Development), placed on titanium alloy inserts within a six-well plate, and maintained in 2.5 mL of KSFM supplemented with 5% (v/v) charcoal-stripped FBS and 50 nM R1881 (PerkinElmer) ^25^. Slices were cultured overnight, rotating at a 30° angle, at 37°C with 5% CO_2_. Alternating slices were collected for RNA extraction and formalin fixation. For gene expression analysis, RNA isolation and RT–qPCR were performed as described above. Only slices with confirmed benign epithelial content (high expression of KRT8 and undetectable *PCA3* by RT–qPCR) were included in the analyses.

### Immunohistochemistry of tissue slice explants

Formalin-fixed paraffin-embedded slices were sectioned into 5 μm sections, deparaffinized, processed for steam antigen retrieval (10 mM sodium citrate, 0.05% Tween 20, pH 6), and stained with anti-megalin (1:500; ab76969, Abcam) overnight at 4°C. A rabbit secondary antibody HRP/DAB kit was used for visualization with hematoxylin counterstaining (ab64261, Abcam).

### Tissue microarray (TMA) immunofluorescent staining and analysis

The TMA contained 118 prostate biopsy cores from 29 patients (20 AA, 9 EA) and at least two benign and cancer cores from each patient. A board-certified pathologist reviewed each core to confirm the cancer grade mark regions for exclusion if they contained artifacts or benign areas intermixed with cancer. Sections (5 μm) were incubated with rabbit polyclonal anti-megalin (ab76969) diluted 1:100 and mouse monoclonal anti-panCK (AE1/AE3) diluted 1:2,000, followed by incubation with secondary antibodies Alexa Fluor 488 goat anti-rabbit diluted 1:200 and Alexa Fluor 555 goat anti-mouse diluted 1:200 (Life Technologies, Carlsbad, CA, USA) and counterstaining with DAPI. Sections were scanned at ×20 on a Vectra3 multispectral imaging system (Akoya Biosciences, Marlborough, MA, USA). Epithelial areas were identified and segmented by machine learning using the panCK marker and HALO software (Indica Labs, Albuquerque, NM, USA) and adjusted manually to ensure accuracy. Epithelial megalin fluorescence intensity was quantified and reported as the average intensity per pixel of the segmented area of each core using the Inform software. The Mann–Whitney U test was used to compare benign cores to PCa cores for all men, EA only, and AA only.

### *LRP2* expression in public datasets

RNA-sequencing datasets for PCa tumors were identified and data exported from cBioPortal^26^. *LRP2* expression was analyzed using analysis of variance (ANOVA) with Kruskal-Wallis for multiple comparisons.

### Statistics

The statistical analysis methods used in each experiment are detailed in the figure legends and methods section.

### Data availability

All data generated or analyzed during this study are included in this published article (and its supplementary information files).

## RESULTS

### Prostate cells express megalin and import SHBG-bound T

Since we previously detected megalin in prostate tissues^21^, megalin transcript (*LRP2*) and protein expression were quantified in prostate cell lines. Both genes and proteins were detected in benign primary patient-derived prostate epithelial cells (PrE AA1 and PrE AA2), immortalized benign prostate epithelial cells (957E-hTERT), and PCa cell lines (LNCaP and 22Rv1) (**Figure 1A**). All cell types expressed the vitamin D receptor (VDR); however, benign PrE cells (AA1, AA2, and 957E-hTERT) lacked androgen receptor (AR) expression (**Figure 1A**). PCa cells (LNCaP and 22Rv1) showed low to no expression of CYP27B1 (**Figure 1A**) and were unable to metabolize 25D to the active hormone 1,25D, as shown by the lack of CYP24A1 induction (**Figure S1**). LNCaP and 22Rv1 PCa cell lines were used to examine testosterone (T) import and AR activity in vitro, as they have differential expression of megalin and AR. When treated with T alone and SHBG-T, both LNCaP and 22Rv1 cells showed AR activation, as evidenced by the increased *KLK3* mRNA levels (**Figure 1B**). SHBG was added in 20-fold excess and incubated with T for 30 min before treating cells to ensure thorough hormone-globulin binding. To inhibit megalin, the cells were pre-incubated with receptor-associated protein (MEG-Inh)^27^(**Figure 1B**). Both LNCAP and 22Rv1 cells exhibited decreased *KLK3* gene expression and ARE luciferase when cells were pretreated with MEG-Inh before hormone treatment; however, the magnitude of inhibition was higher in 22Rv1 cells, which expressed more megalin. Pre-incubation with MEG-Inh did not block the response to added T alone, demonstrating its specificity to SHBG-T. To visualize SHBG/T internalization, we used Alexa Fluor 555-labeled human SHBG (SHBG-555). SHBG-555 was localized to the plasma membrane of LNCaP and 22Rv1 cells in response to SHBG and SHBG + T treatment (**Figure 1C**). The addition of SHBG-T resulted in greater internalization and punctate patterns than SHBG treatment alone, and MEG-Inh blocked the internalization of SHBG. These data demonstrate that SHBG-bound T cells are present in prostate cells in a megalin-dependent manner.

**Figure 1.**
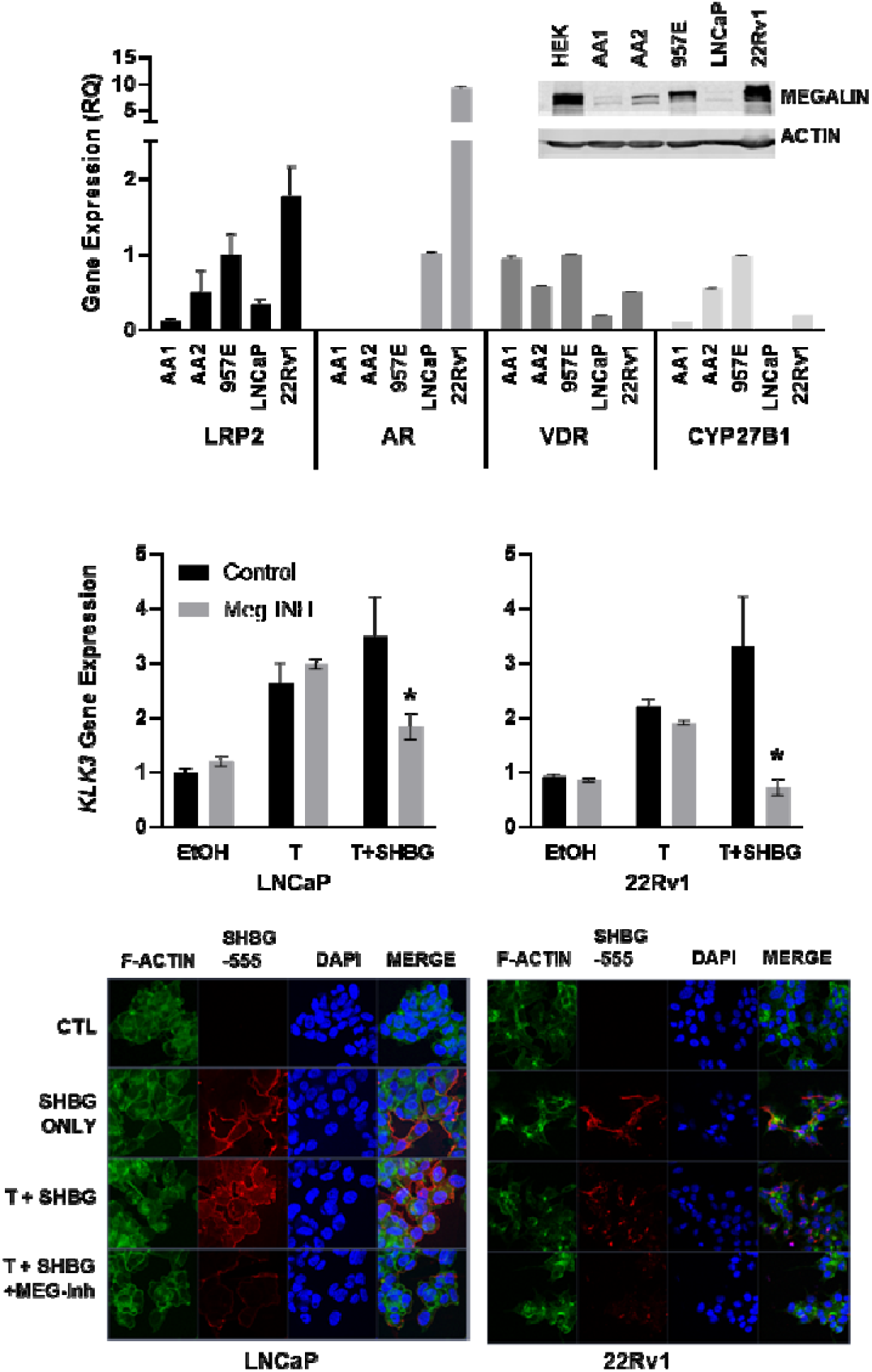
Prostate cells express megalin and import SHBG-bound T. (A) Gene expression of *LRP2, AR, VDR* and *CYP27B1* in a panel of prostate cell lines as shown by RT-qPCR shown as relative quantitation to *HPRT1.* Error bars are standard error of the mean (SEM). (A, inset) Western blot for megalin in prostate cell line panel. (B) Regulation of *KLK3* (prostate-specific antigen [PSA] gene) expression after 24 h following treatment with vehicle control (CTL), 50 nM T alone (T), 50 nM T preincubated with 500 nM SHBG (SHBG), or T + SHBG in cells preincubated with 1μM MEG-Inh in LNCaP and 22Rv1 PCa lines. (C) Visualization (×63) of SHBG-555 (red) import into cells with DAPI (blue) nuclear and F-actin (green) cytoskeletal counterstains. Statistical analysis was performed using a one-way ANOVA with a two-stage linear step-up procedure of Benjamini, Krieger, and Yekutieli for multiple comparisons; \**P* < 0.05 for comparison to CTL, All graphs represent mean ± SEM of three or more individual experiments with two replicates per experiment.

### Knockout of *Lrp2* in mouse prostate reduces T

To determine whether prostate T levels are affected by the absence of megalin, we created a prostate-specific knockout of *Lrp2*, since obligate *Lrp2* knockout causes perinatal lethality in most affected animals ^23,28^. A tamoxifen (TAM)-inducible prostate-specific knockout of *Lrp2* was generated by crossing an *Lrp2*-floxed mouse^29^ with a probasin-driven TAM-inducible Cre recombinase mouse (Pb-MerCreMer)^30^ (**Figure 2A**). No breeding problems were encountered with the homozygous bitransgenic line. *Lrp2*^fl/fl^/*Cre*^+/+^ mice and control (*Lrp2*^fl/fl^ only and *Cre^+/+^* only) mice were injected with TAM at postnatal day 10 (P10-TAM), which resulted in recombination of genomic DNA only in the prostate and not in mouse tails (**Figure 2B**). Prostates, testes, and sera were collected at 24 and 32 weeks of age for androgen measurement by liquid chromatography-tandem mass spectrometry (LC-MS/MS). Prostate T and DHT levels were significantly lower in *Lrp2*^fl/fl^/*Cre*^+/+^ mice than in the control mice (**Figure 2C**). Serum T and testes T levels were tightly correlated in all mice, supporting testes as the source of serum T. However, neither prostate T nor DHT was significantly correlated with serum T in control and *Lrp2*^fl/fl^/*Cre*^+/+^ mice (**Figure 2D**), suggesting that prostate T levels are not due to passive diffusion from the serum. In addition, prostate T and DHT were only significantly correlated in the control *Lrp^f^* group, suggesting that the regulation of T to DHT is different in prostates lacking *Lrp2*. Serum DHT was undetectable in most mice, consistent with T as the primary circulating hormone. Although the differences in prostate androgens were significant, there were no differences in prostate histology, prostate weight, or fertility (data not shown). The relationship between serum and tissue hormone levels supports a regulated transport mechanism of T into the prostate, rather than passive diffusion of serum T.

**Figure 2.**
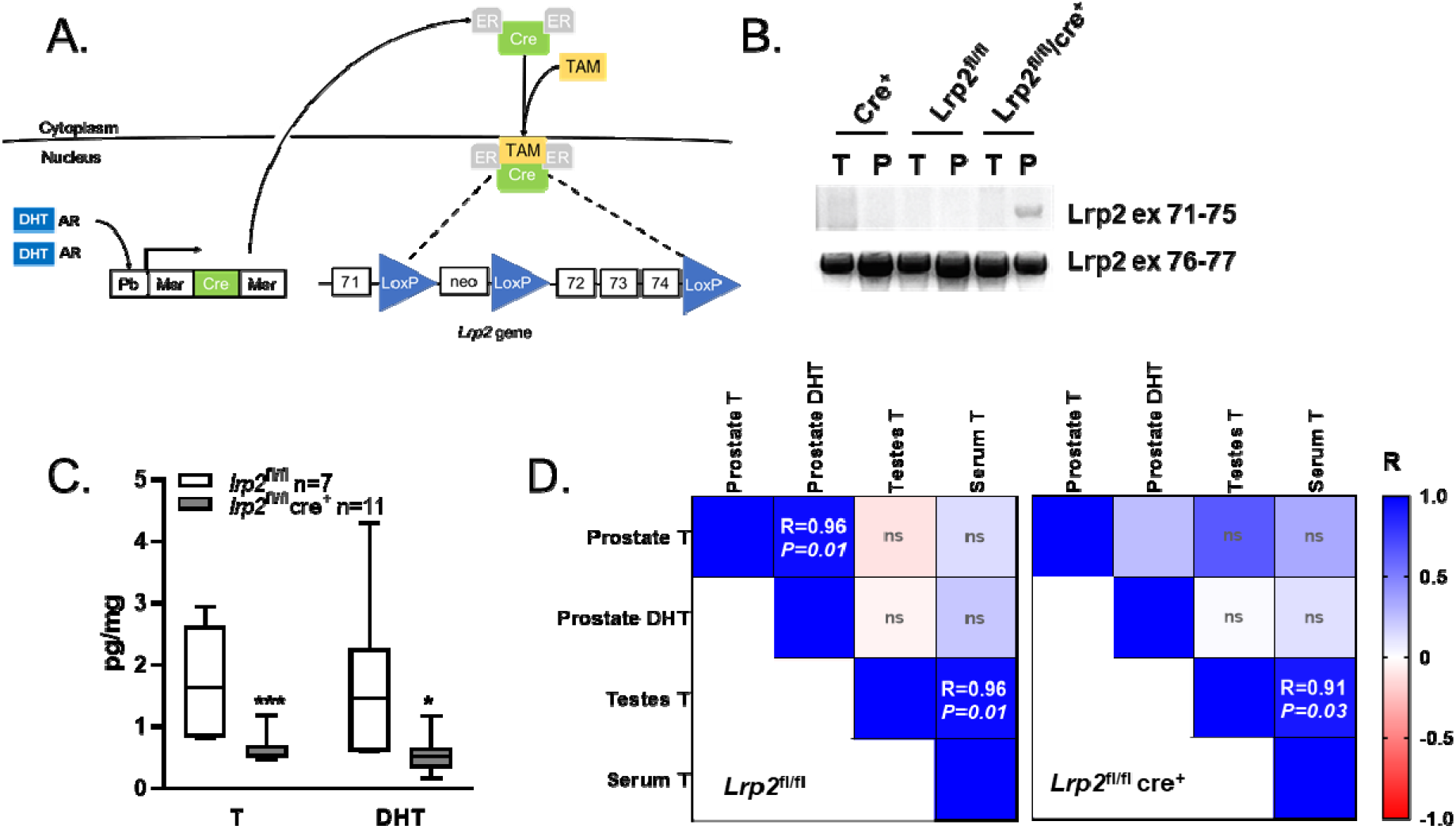
Loss of *Lrp2* in the mouse prostate reduces prostatic androgen levels. (A) Mouse model of conditional knockout of *Lrp2* in prostate epithelium. (B) Recombination of *Lrp2* exons 71–75 after TAM treatment in prostate DNA (P), but not in tail DNA (T), only in bitransgenic mice. PCR for exons 76–77 was used as the positive control. (C) Prostate levels of T and DHT quantified by LC-MS/MS for *Lrp2*^fl/fl^ and *Lrp2*^fl/fl^/*cre*^+/+^. The graphs show the mean with maximum-minimum bars. (D) Heat map of Pearson correlation coefficients (*R*) for tissue and serum androgens in P10-TAM mice (*Lrp2*^fl/fl^ n=5, *Lrp2*^fl/fl^/*cre*^+/+^n=6). The *P*-value is shown within each cell; ns, not significant.

### Vitamin D negatively regulates *LRP2* expression

We previously observed a negative correlation between serum 25D levels and *LRP2* prostatic expression in AA men^21^, suggesting that *LRP2* is regulated by 25D. Therefore, we sought to characterize the effect of 25D on *LRP2* gene and megalin protein expression in vitro. Primary prostate epithelial cells (PrE-AA1) treated with 25D exhibited decreased *LRP2* gene and megalin protein expression (**Figure 3A-B**). To assess the regulation of *LRP2* expression at the transcriptional level, we characterized the *LRP2* promoter activity in vitro. The *LRP2* promoter was cloned into the *Renilla* luciferase reporter plasmid (*LRP2-Rluc*), which was suppressed by 25D in 957E-hTERT cells (**Figure 3C**). 957E-hTERT cells were used for luciferase experiments because they can be transfected at high efficiency, whereas PrE cells are difficult to transfect. Furthermore, 957E-hTERT cells are immortalized prostate cells that do not express AR and are phenotypically similar to PrE cells^31^. VDR forms an obligate heterodimer with the retinoid X receptor (RXR) and binds to the vitamin D response elements (VDREs). Analysis of the *LRP2* promoter fragment identified multiple RXR:VDR motifs (**Figure 3D**; **Figure S2**). (**Figure 3C**). These transcriptional analyses support the hypothesis that *LRP2* expression is regulated by hormone-stimulated transcription factors in response to 25D.

**Figure 3.**
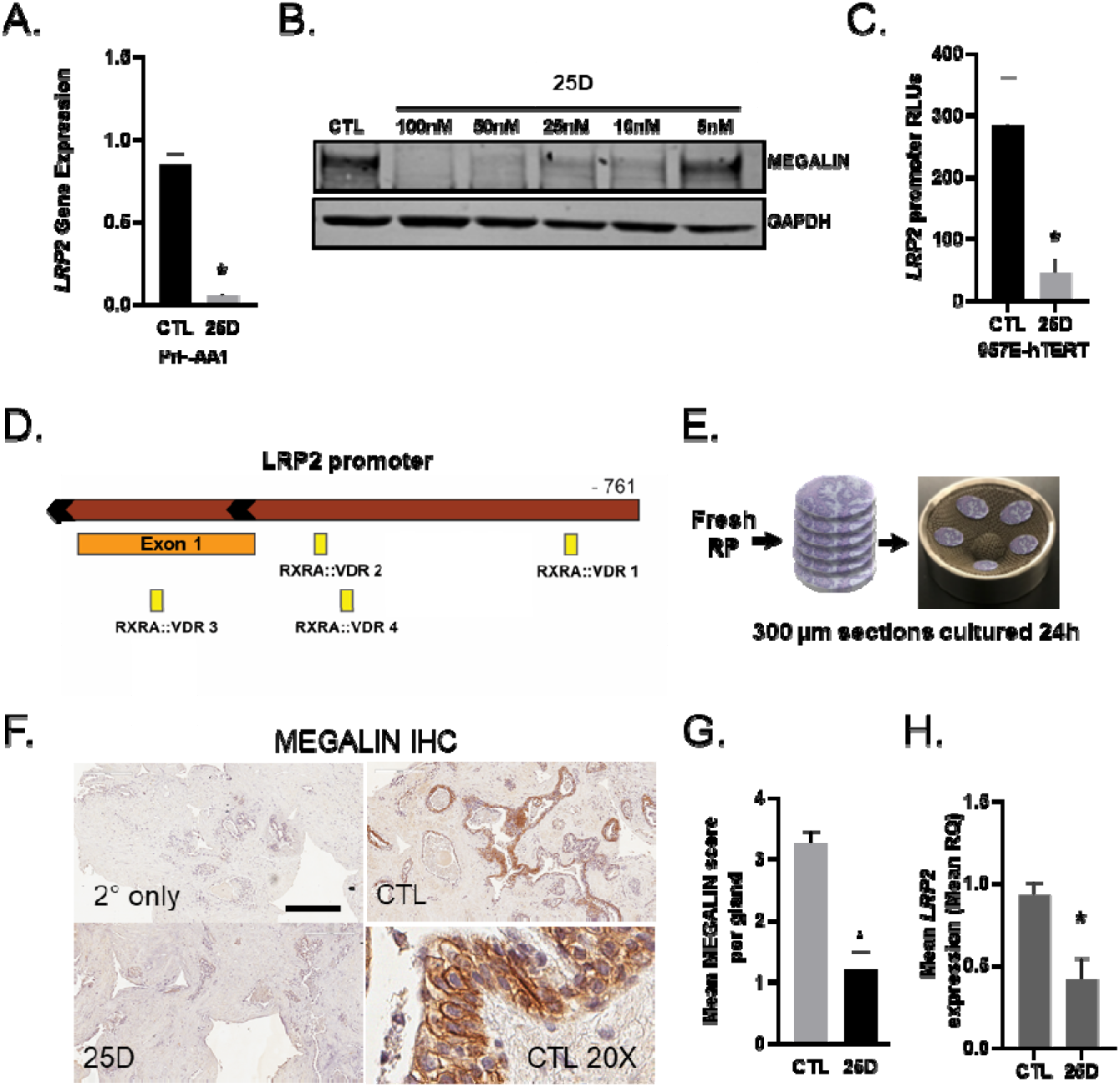
Negative regulation of *LRP2* and megalin proteins by vitamin D in prostate cells and tissue slice explants. (A) *LRP2* expression following 24 h of treatment with 50 nM 25D in PrE-AA1 cells. (B) Immunoblot for megalin after 48 h of 25D treatment of PrE-AA1 cells. (C) Activity of a custom *LRP2* promoter luciferase construct after 24 h of 50 nM 25D treatment in 975E-hTERT cells; RLU, relative luciferase units normalized to the transfection control. (D) *LRP2* promoter contains the RXR:VDR, and AR-binding motifs. (E) Ex vivo prostate tissue slice workflow. (F) Images and quantification of (G) megalin protein and (H) LRP2 gene expression by IHC in tissue slices after 24 h of treatment with 50 nM 25D. Scale bar = 150 μm. The graph shows the mean pathologist score per gland. Graphs represent the mean ± SEM from triplicate experiments. For tissue slices, graphs show the representative experimental mean ± standard deviation (SD) with two replicates per experiment. *P* values were determined using an unpaired *t*-test.

PrE and 957E-hTERT cells respond to 25D but did not express AR. Thus, we examined these responses in fresh ex vivo benign human prostate tissue slice cultures (**Figure 3E**), which express all components of androgen and vitamin D activation/response pathways, including *CYP27B1* (vitamin D 25-hydroxylase), *VDR, LRP2, SRD5A* (T to DHT conversion), and *AR* (data not shown). Hormone responsiveness was demonstrated by the robust induction of *CYP24A1* and *KLK3* gene expression by 25D and T, respectively, alone or in the presence of their serum-binding globulins, DBP, and SHBG (**Figure S3**). Megalin protein and *LRP2* expression were decreased in tissue slices treated with 25D (**Figures 3F-H**), consistent with our observations in cell lines and the relationships we previously observed between serum 25D and prostate *LRP2* in patient data^14^. These findings strongly support a vitamin D-mediated negative feedback on *LRP2* expression.

### Intraprostatic DHT is higher in AA men and inversely correlates with vitamin D status

Negative regulation of megalin by vitamin D suggests that vitamin D deficiency may lead to megalin upregulation and, subsequently, increased prostate import of steroid hormones, including SHBG-bound T. To test this hypothesis, we examined the relationship between these hormones in patients. We quantified DHT in frozen radical prostatectomy tissues from a cohort of PCa patients for whom we had previously measured vitamin D metabolites^14^. Vitamin D status, as measured by serum 25D level, was negatively correlated with intraprostatic DHT (**Figure 4A**) in all patients but was not significant when analyzed separately by ancestry. In this cohort, AA patients had higher prostate levels of the active hormone DHT than EA men (**Figure 4B**). DHT was the predominant androgen in the prostate, whereas T levels were undetectable in the majority of patients (**Figure 4B**), supporting metabolism from T to DHT once in the tissue. In the serum, T was the dominant androgen and was slightly lower in AA men than in their EA counterparts, which was similar to their 25D status (**Figure 4C**).

**Figure 4.**
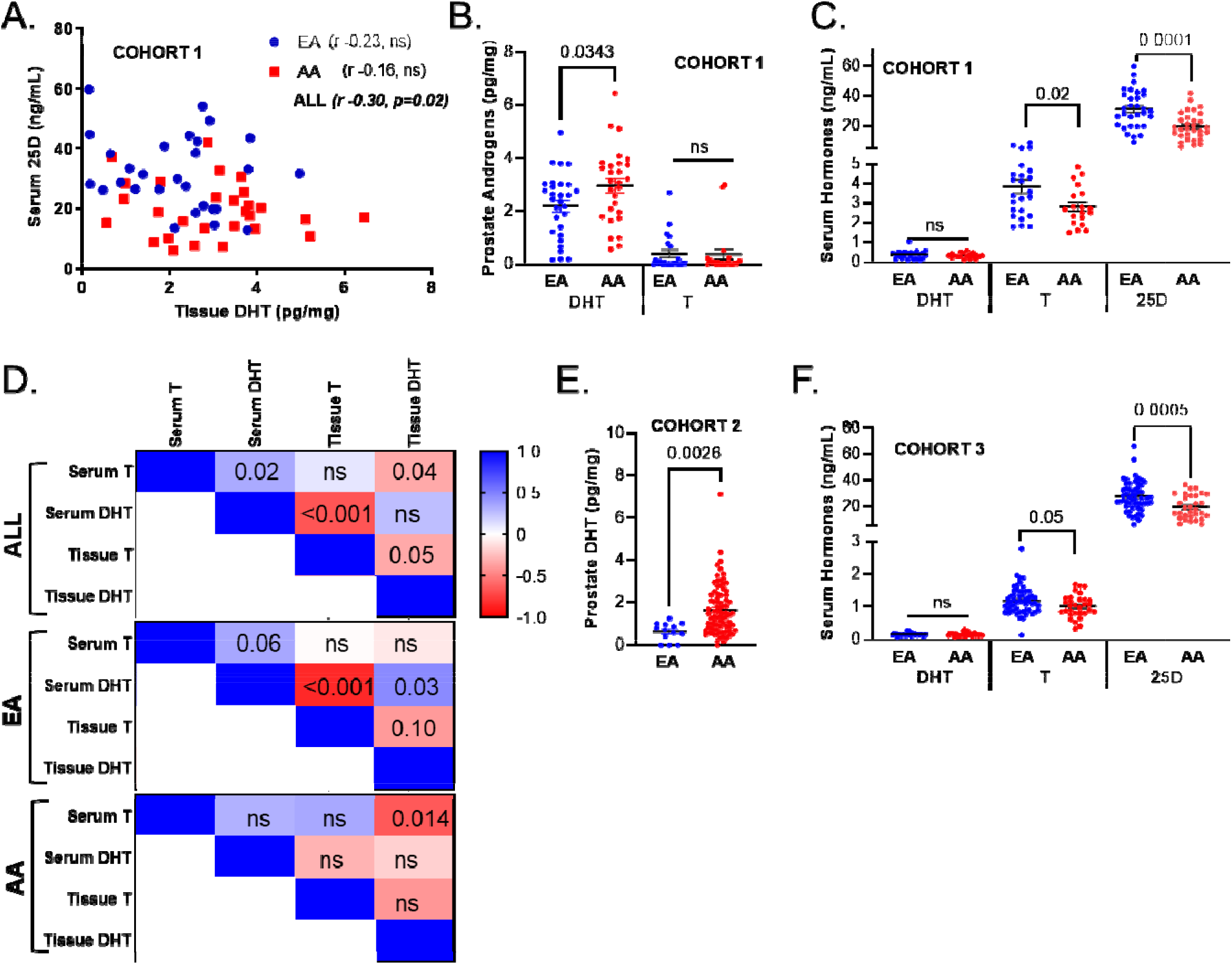
Ancestry-specific differences in androgen levels and relationships between DHT and 25D. (A) Correlation between serum 25D and prostate DHT in EA (*n* = 29) and AA (*n* = 28) men in Cohort 1. Correlation values (*r*) were determined using Spearman’s rank test. (B) Prostate tissue levels of T and DHT in Cohort 1. (C) Serum levels of T, DHT and 25D in Cohort 1. (D) Spearman correlations (R shown in heatmap) between serum and prostate tissue concentrations of hormones. P values shown within cells. (E) Prostate tissue levels of DHT in Cohort 2 (EA *n*=13, AA *n*=82). (F) Serum levels of DHT, T and 25D in Cohort 3 (EA *n*=33, AA *n*=28). All hormone measurements by LC-MS-MS. Graphs represent mean with 95% confidence interval (95% CI). The *P* values were determined by unpaired two-tailed *t* test.

The free hormone hypothesis would be supported by the positive correlations between serum and tissue androgen levels, but this is not what we observed. Correlation analyses between prostate tissue and serum androgen levels showed that only EA men had a positive correlation between serum DHT and tissue T (**Figure 4D**). In contrast, an inverse correlation was observed between serum T and prostate tissue DHT in the overall cohort and in AA men (**Figure 4D**). There was a very high correlation between serum DHT and tissue T in EA men only, which may be due to the low levels of both in the samples (**Figure 4D**).

We measured prostate DHT in a second cohort of patients with PCa and observed a similar difference in tissue DHT between the AA and EA groups (**Figure 4E**). However, there were very few EA men in this cohort, and we did not have matched serum. In a third cohort from PCBN, serum levels of hormones were similar to those in cohort 1, with lower T and 25D in AA men than those of white men(**Figure 4F**).

Overall, the inverse correlation between serum 25D and intraprostatic DHT supports our hypothesis that serum vitamin D deficiency drives higher prostatic DHT levels. Second, the relationship between hormones in the serum and tissues differs by ancestry and does not support the free-hormone hypothesis in AA men.

### Megalin and *LRP2* levels are lower in PCa

A tissue microarray (TMA) composed of prostate cores from 29 patients (20 AA, 9 EA), with four cores from each patient, two benign and two PCa regions per patient, was stained for megalin. Fluorescence intensity was digitally quantified in the epithelial regions using the epithelial marker Pan-CK to segment the epithelium (**Figure 5A**). Neither frozen tissues nor serum were available for patients represented in this TMA; therefore, we could not correlate the hormone levels with megalin expression. However, megalin protein levels were significantly lower in PCa tissues than in benign tissues (**Figure 5A**). The majority of the PCa on this TMA was Gleason 3, with only five cases of Gleason 4, and no difference by Gleason was observed in this small cohort.

**Figure 5.**
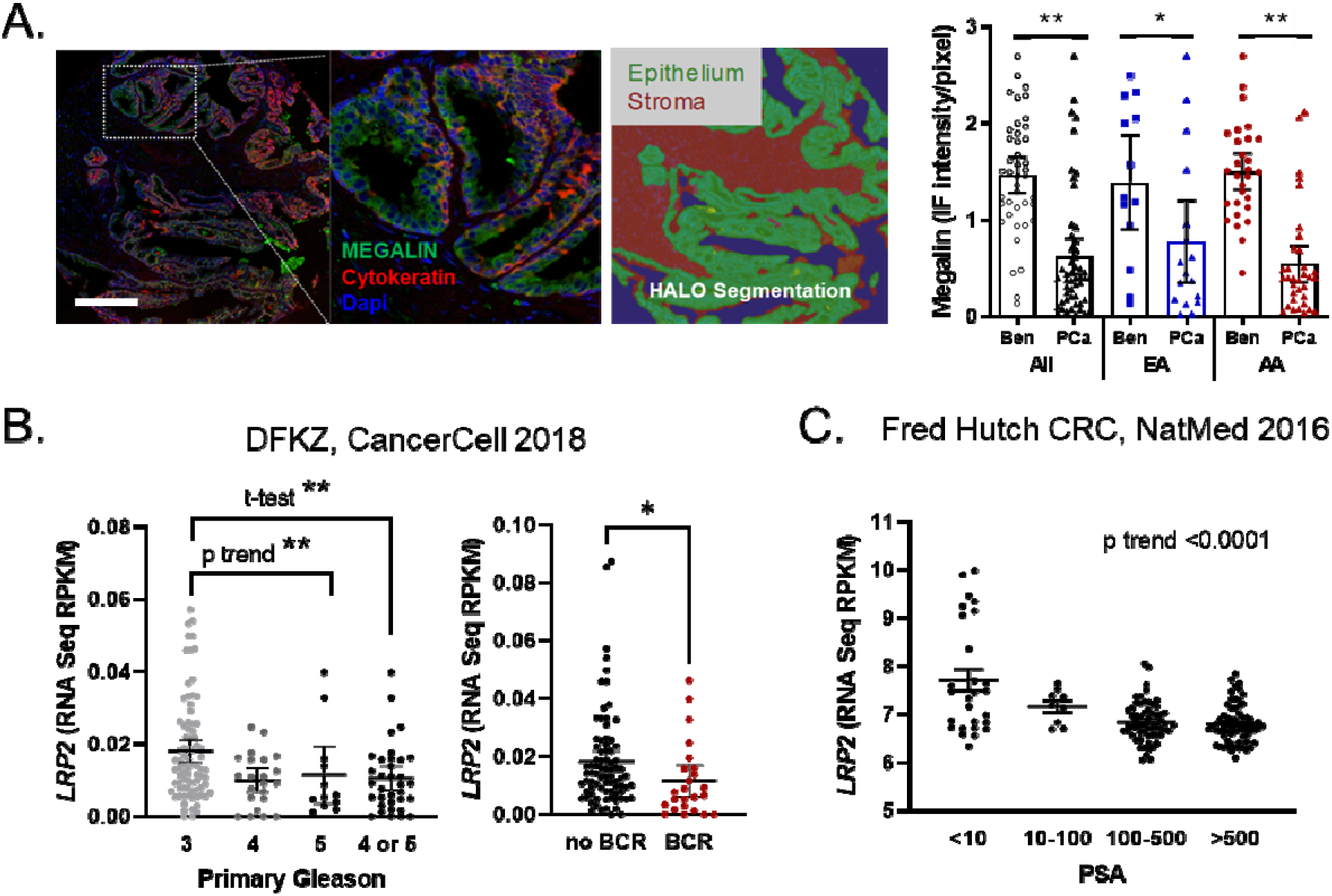
Megalin expression is dysregulated in cancer and lower with disease aggressiveness. (A) Digital quantitation of epithelial megalin expression on a TMA consisting of 118 prostate biopsy cores from 29 patients (EA, *n* = 9; AA, *n* = 20). Epithelial regions were segmented by panCK staining, and benign and cancer regions were determined by a board-certified pathologist. Scale bar = 100 μm. Immunofluorescence (IF) intensity per pixel for megalin expressed as mean ± SEM; Ben, benign. (B) *LRP2* expression in RNAseq data set of primary PCa tumors by primary Gleason grade and future BCR from Gerhauser et al. Analyses by Gleason was ANOVA with post test for linear trend and unpaired t test between Gleason 3 and 4 or 5 combined. BCR *P* values were determined by a Mann-Whitney test. \**P* < 0.01.**P<0.001 (C) *LRP2* expression in RNAseq data set of castration resistant PCa tumors by serum PSA from Kumar et al^33^. ANOVA followed by post test for linear trend.

*LRP2* expression was assessed in larger cohorts, using publicly available datasets. Analyses of gene expression data from a Dana Farber cohort of localized PCa^32^ showed that *LRP2* levels decreased with primary Gleason grade (**Figure 5B, left panel**) and were lower in patients who had future biochemical recurrence (BCR) (**Figure 5B, right panel**). In a cohort of castration-resistant tumors in 63 men, PCa *LRP2* levels were the highest in men with the lowest PSA levels in the cohort^33^ (**Figure 5C**). Together, the PCa data support that *LRP2* expression is highest in normal tissues that are most differentiated, which is consistent with the localization of megalin in mature luminal cells.

## DISCUSSION

This study follows our recent finding that megalin protein is expressed in the membrane of the prostate epithelium and that *LRP2* gene expression correlates with vitamin D status and the percentage of African ancestry^21^. These findings led us to hypothesize that megalin is part of a compensatory pathway that is upregulated during vitamin D deficiency to increase the import of hormones into the prostate. Here, we focused on how this increase in megalin affects the import of SHBG-bound T, which is also known to be imported by megalin. We showed that megalin imports SHBG-bound T, is regulated by vitamin D, and is dysregulated in PCa. These findings provide a direct link between vitamin D deficiency and PCa disparities in AA men.

Given the pronounced role of androgens in PCa risk and progression, we examined megalin as a mediator of prostate androgen levels. T is the principal circulating androgen with 70% bound to SHBG, approximately 5% free T and the remaining bound to albumin ^17^. T is thought to follow the free-hormone hypothesis, with only free or albumin-bound T available to tissues. We showed that megalin binds and internalizes SHBG-bound T, which is similar to other reports in LNCaP cells ^17^. This was observed in two PCa cell lines, in which SHBG-bound T was internalized, resulting in *KLK3* induction. We further demonstrated that the loss of megalin in mouse prostate epithelium decreased prostate androgen levels. Although knockout mice demonstrate some regulation of prostate T by megalin, it is important to note that mice differ from humans in that mice primarily circulate albumin-bound T rather than SHBG-T. Megalin is a multiligand receptor that also mediates albumin uptake ^34^. Our findings do not exclude the presence of other SHBG-T uptake receptors ^35^. The data shown here support megalin-dependent import of SHBG-T into the prostate and are consistent with full *Lrp2* knockout^19^, which displays impaired descent of the testes into the scrotum, and other defects consistent with sex steroid disruption.

A negative feedback loop was previously reported for vitamin D skeletal myotube cultures, in which growth at high levels of 25D significantly decreased DBP-bound 25D uptake in cultures ^36^, although megalin was not specifically implicated in that report. We also observed that high levels of 25D decreased the expression of *LRP2* and megalin proteins. We acknowledge that our findings present a paradox in that *LRP2* expression is regulated by the extracellular amounts of hormones, whereas we observed higher levels of androgens and vitamin D metabolites within the prostates of AA men. It was unexpected that the serum levels of hormones would correlate with *LRP2* expression rather than tissue levels. These findings reveal the complexities of endocrine hormone regulation within tissues and suggest there may be a role of membrane vitamin D receptor, which has been previously reported in other contexts^37,38^. These findings not only support a preventive role for vitamin D but also the need to avoid vitamin D deficiency in PCa patients treated with anti-androgen therapies.

The presence of a compensatory feedback loop to regulate intraprostatic hormone levels has implications for the AA population, who are disproportionately vitamin D-deficient ^39^. We analyzed vitamin D metabolites in three cohorts of patient specimens. We had both serum and tissue samples from Cohort 1 and found that serum 25D and prostatic DHT were significantly inversely correlated, consistent with vitamin D regulation of hormone import via megalin. Moreover, AA men had higher levels of DHT in prostate tissue than EA men in cohort 1 and in a second cohort (2) with only prostate tissues. Elevated prostatic DHT may directly contribute to the increased incidence of aggressive PCa and PCa mortality among AA men as lower prostate androgens have been shown to decrease the risk of PCa and PCa mortality^40–42^. AA men also presented with lower levels of serum T in cohort 1 and cohort 3 (serum only), further supporting the active transport of SHBG-T into the prostate rather than passive diffusion of T. This relationship outlines an intricate, yet detrimental interaction between the androgen and vitamin D axes that characterizes the adverse effects of a double disparity in men of West African descent.

Our serum finding that serum T is lower in AA men differs from previous studies that found no racial differences in serum total T or free T ^43,44^. Hu et al. reported a more rapid age-related decline in serum T in AA men than in white men^45^, although they did not observe lower levels in AA men. This discrepancy may reflect differences in assays, sample preservation methods, and cohort sizes. We also showed that the relationship between serum and prostate tissue androgen levels differed between AA and EA. Although our results were significant, we acknowledge that there is a limitation in the under-representation of vitamin D-replete AA patients and vitamin D-deficient EA patients in the cohort, which is needed to separate ancestry from deficiency.

Given the dependence of PCa on androgens and the prior report of LRP2 polymorphisms with PCa aggressiveness^20^, we examined megalin protein and *LRP2* gene expression in multiple cohorts of patients with PCa. Megalin protein levels were markedly lower in cancer tissues than in benign tissues from radical prostatectomy samples. In publicly available cohorts, *LRP2* expression is further decreased by Gleason, which increases with a lack of differentiation, suggesting that its expression is connected to differentiated cells. We also found that patients who had BCR had lower *LRP2*, which is likely related to its relationship with Gleason score. These findings suggest that PCa may be less dependent on megalin and utilize free hormones once established.

An important consideration regarding race-related differences is that race is a social construct and proxy for ancestry. We do not suggest that biological differences exist because of patient’s self-declared race. Rather, because vitamin D status is directly correlated with skin pigmentation, our findings suggest that vitamin D supplementation may reduce the levels of prostate androgens, which would mostly affect AA men who tend to be vitamin D deficient.

In conclusion, our in vitro and ex vivo data show that megalin is functional in the prostate, transporting SHBG-bound T into the cell, which complicates the free hormone hypothesis. We also show that megalin expression is negatively regulated by vitamin D, and, in times of deficiency, megalin is upregulated, potentially increasing the import of both vitamin D and T. This may signify a once-protective compensatory mechanism of vitamin D gone awry, which increases the likelihood of androgen import and increases the probability of harmful androgen actions that may contribute to the disparity in PCa aggressiveness that plagues AA men.

## ACKNOWLEDGEMENTS

Tissue samples were provided by the UI Health Biorepository, NCI Cooperative Human Tissue Network (CHTN), and Prostate Cancer Biorepository Network (PCBN), which was supported by the Department of Defense Prostate Cancer Research Program. We thank D. Klara Valyi-Nagy and Alexandru Cristian Susma from the UI Health Biorepository, Ryan Deaton, and the UIC Research Histology and Tissue Imaging Core for assistance with the tissue microarray analysis, Vicky Macias and Andre Kajdacsy-Balla for assessment of pathology of patient tissues, Morgan Zenner and Michael Schlicht for assistance with generating tissues slices, and Hui Chen for editing the methods. Finally, we thank the patient participants who donated their specimens for research and Drs. Michael Abern, Daniel Moreira, and Simone Crivellaro for the procurement of radical prostatectomy tissue specimens.

## Supplemental Figures

**Figure S1.**
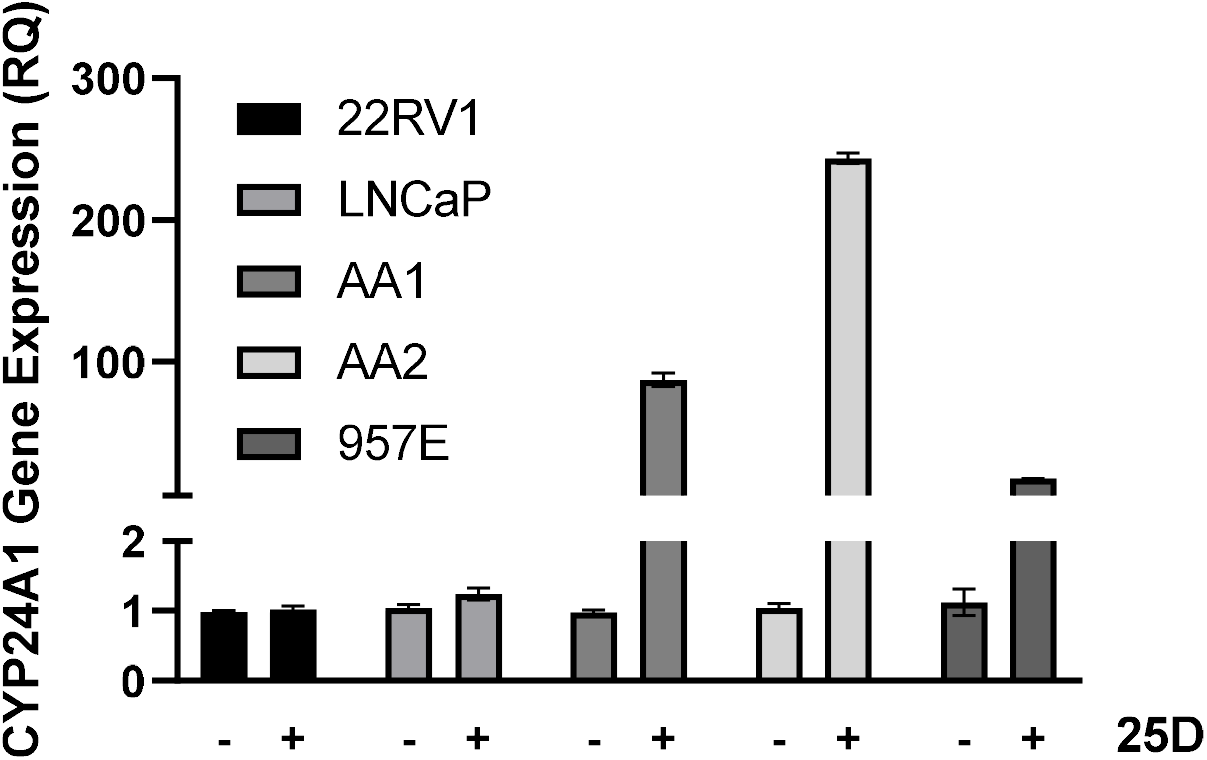
Accompanies Figure 2. **A,** PCa cells lines do not respond to 25D. RT-qPCR for *CYP24A1* following 16 hours of 50 nM 25D treatment in serum free conditions. **B**, Megalin inhibitor does not impact activity and import of T alone. RT-qPCR for *KLK3* following 16 hours of 10 nM T treatment. Cells were pretreated with 1 μM MEG-inhibitor for 1 hour before T. Expression shown as relative quantitation to *HRPT.* Error bars are SEM. *p<0.01

**Figure S2.**
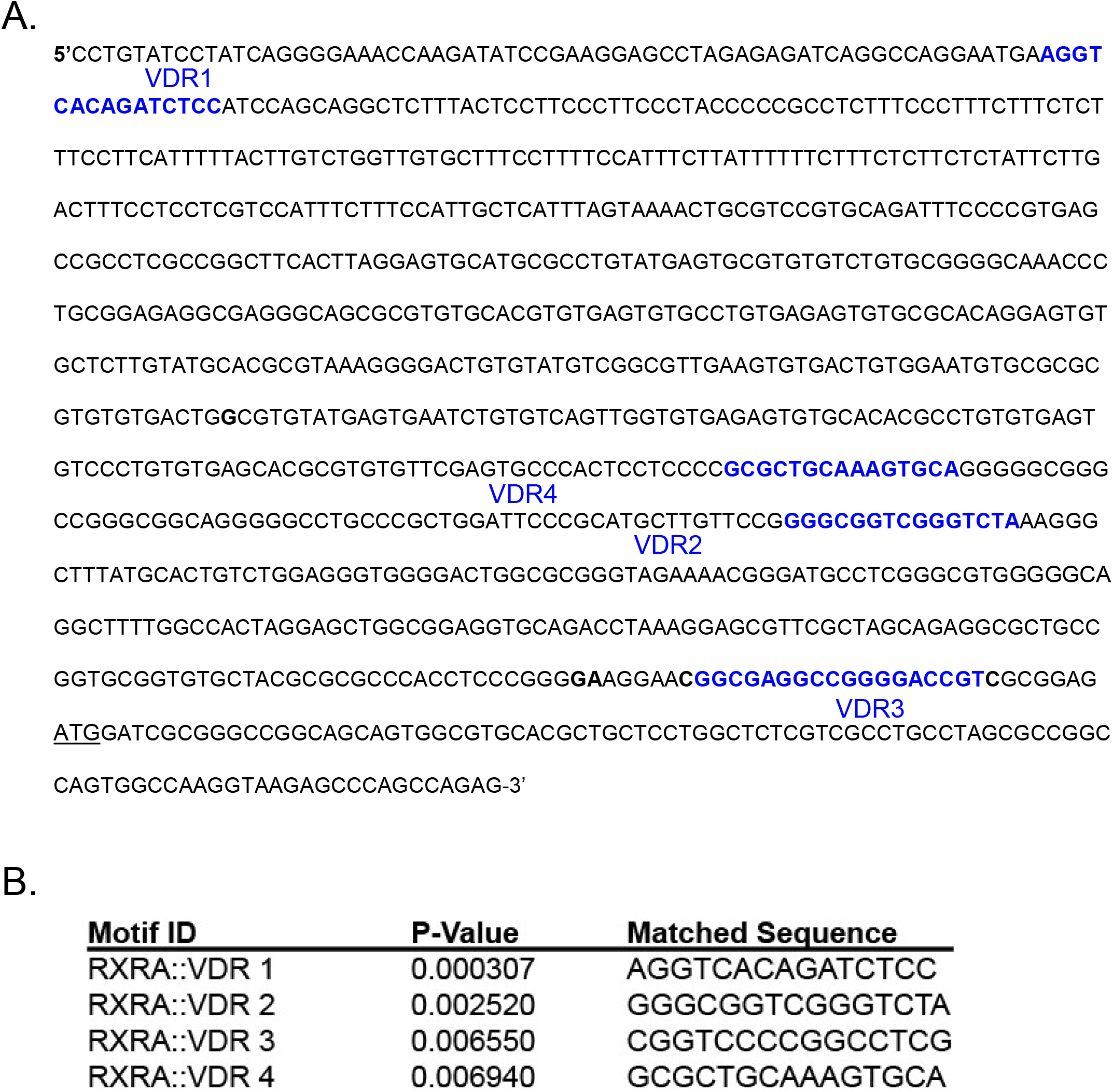
Vitamin D receptor and androgen receptor response elements in LRP2 promoter. **A,**-791 bp of LRP2 promoter showing mapped areas for VDR and AR response elements. **B,** Jasper prediction of the binding sites with P-value. Note that VDR binds VDREs as an obligate heterodimer with RXRα.

**Figure S3.**
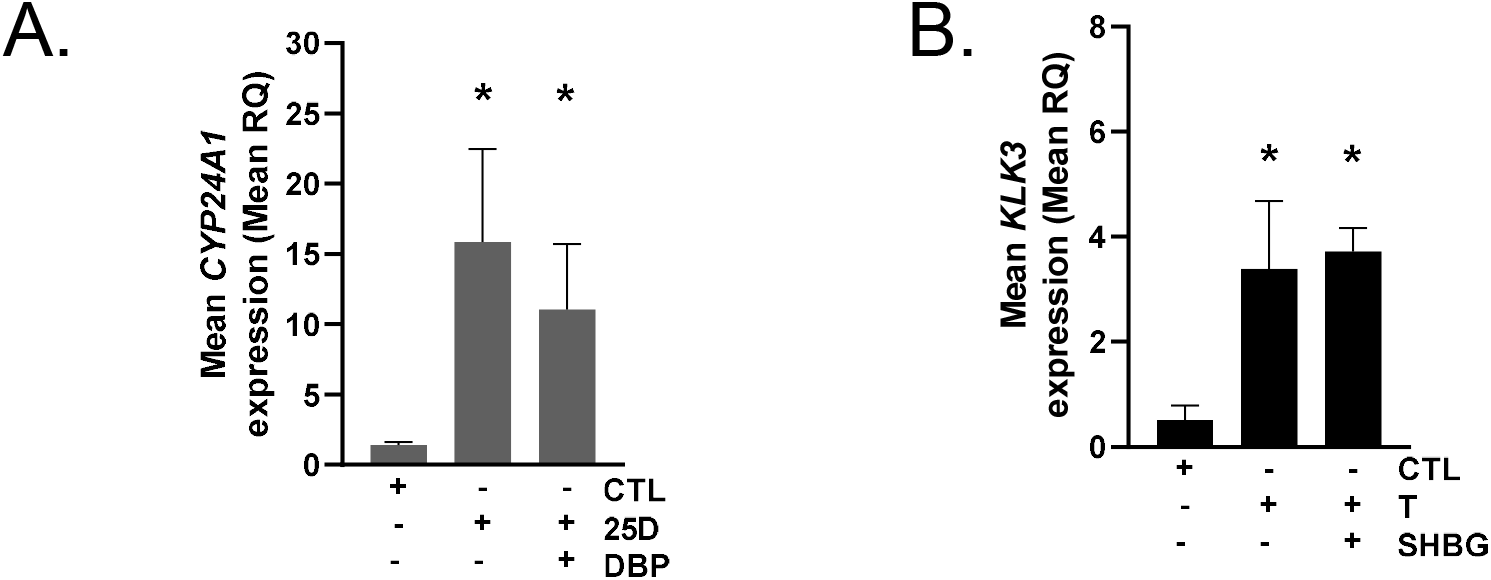
Prostate slices respond to 25D and T. **A,** Expression of *CYP24A1*, a vitamin D response gene, in slices treated for 24 h with 50nM 25D with and without DBP. B, Expression of *KLK3*, an androgen response gene, in slices treated for 24 h with 50nM T with and without SHBG. Data shown as mean and SEM of biological replicates from 3 patients.

**Table S1.** Cell and tissue characteristics.

| **MODEL** | **TISSUE** | **MORPHOLOGY** | **DISEASE** | **ETHNICITY** |
| --- | --- | --- | --- | --- |
| **PrE-AA1** | Prostate | Epithelial | Benign | African American |
| **PrE-AA2** | Prostate | Epithelial | Benign | African American |
| **957E-hTERT** | Prostate | Epithelial | Cancer (local) | White European descent |
| **HEK293** | Kidney | Epithelial | Benign | Unknown |
| **22Rv1** | Prostate | Epithelial | Malignant origin<br>Behaves benign | White European descent |
| **LNCaP** | Prostate | Epithelial | Cancer (lymph node) | Unknown |
| **TS-AA1** | Prostate | Epithelial | Benign | African American |
| **TS-H1** | Prostate | Epithelial | Benign | Hispanic |

**Table S2.**
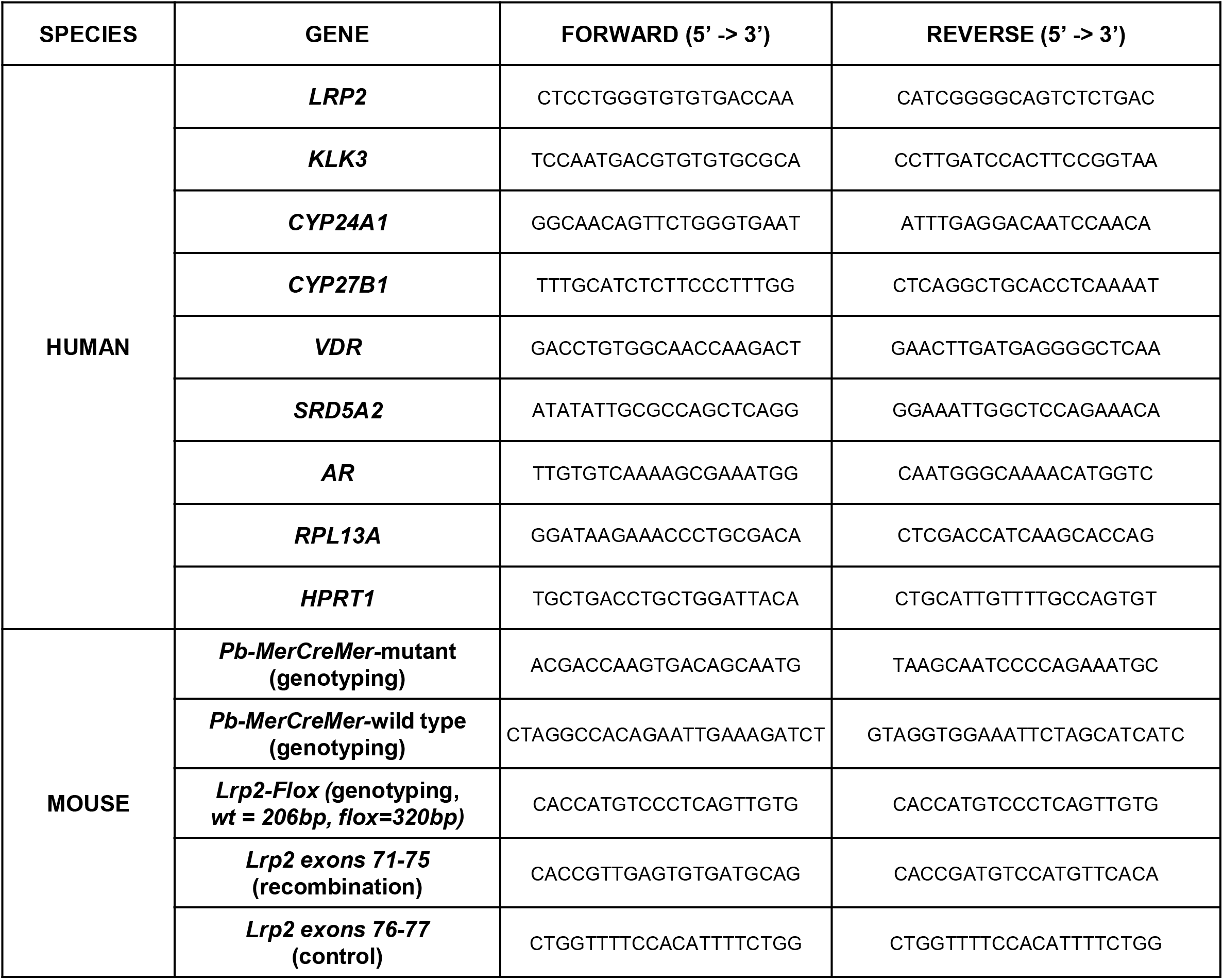
Primer sequences.

